# Medical Xenophobia: The Voices of Women Refugees in Durban, Kwazulu-Natal, South Africa

**DOI:** 10.1101/603753

**Authors:** Yvonne Munyaneza, Euphemia Mbali Mhlongo

## Abstract

**Background:** Women refugees are mostly affected due to their specific needs for reproductive health services. In their attempt to utilize reproductive health care services, women refugees face medical xenophobia by the health care professionals. Upon their arrival in South Africa, refugee women do not undergo any screening, and this exposes them to health risks making them more prone to all different types of diseases, as many of them are survivors of rape and other acts of sexual violence.

**Objective:** The aim of the study was to describe the voices of women refugees regarding reproductive health services in public health institutions in Durban KwaZulu-Natal

**Methods:** A qualitative, descriptive design was used. Data was collected through face-to-face interviews with eight women refugees living in Durban, KwaZulu-Natal. Thematic content analysis guided the study.

**Results:** Two main themes emerged from the data: negative experiences/challenges, and positive experiences. The negative experiences included medical xenophobia and discrimination, language barrier, unprofessionalism, failure to obtain consent and lack of confidentiality, ill-treatment, financial challenges, internalised fear, religious and cultural domination, the shortage of health personnel and overcrowding of public hospitals. The positive experiences included positive treatment, care and social support.

**Conclusion:** The study concluded that discrimination and medical xenophobia remain a challenge for women refugees seeking reproductive health services in public health institutions in Durban, KwaZulu-Natal.

## Introduction

Women refugees just like every other migrant from around the world face some form of xenophobia in the process of trying to settle in their host countries. Women are mostly affected due to their specific needs for reproductive health services. In their attempt to utilize reproductive health care services, women refugees face medical xenophobia by the health care professionals. In the study carried in South Africa in relation to the access to maternal health care services for refugee women, it was noted that refugee women are more disadvantaged as opposed to other women and they face unfair discrimination in public health care institutions as they are ill-treated by health care workers. As a result, foreign nationals including refugees are reluctant to attend the public health care institutions. Thus, it is important to note that reproductive health care is a crucial element for refugees women. ^[1]^

The refugees in general face medical xenophobia during their encounter with health care workers and language barriers and documentations were found to be the biggest challenges and barriers preventing refugees from accessing and utilising reproductive health care services. The xenophobic acts of violence and hostility towards foreigner nationals mostly refugees, resulted in South Africa being labelled as one of the most xenophobic countries worldwide.^[2]^

It was reported that refugees face a number of challenges when dealing with public hospitals. Those challenges include xenophobic attitude from health care service providers, language barriers and lack of documentations. As a result, refugees are denied access to health care services. ^[3]^ Alfaro-Velcamp ^[4]^, in her study she noted that refugees and migrants are discriminated against and denied health care services based on their nationalities and documentations.

Medical xenophobia and discrimination against refugees go hand in hand in the host country and were identified as the most serious barriers to health care access by refugees in general. Refugee women form the highly vulnerable group among refugees and their health are often overlooked. ^[5]^

When refugees arrive in their host city, many need immediate health care services as a result of long journeys, pre-existing conditions, pregnancies or illness contracted on the way. Despite the protocols in place outlining the rights and the need to provide health services to refugees, many refugees like other migrants experience barriers in access of quality health care services. ^[6]^

However, the experiences of refugee women regarding utilization of reproductive health services in KwaZulu-Natal are unknown. We have no idea of their feelings and the challenges they face when trying to access reproductive health services as there is no documentation of such. There is lack of literature on the experiences of refugee women with regard to their access of reproductive health services in South Africa. To describe the challenges faced by refugee women in utilising reproductive health services, we approached ten women refugees aged between 18 and 49 years, who originated from the DRC, Burundi or Rwanda. These women live in Durban, KwaZulu-Natal, and have sought reproductive health services at one of the public hospitals in the city of Durban

Refugee women from the three Great Lakes countries who were in their reproductive ages, living in Durban and having sought reproductive health services at one of the public health facilities situated in the city of Durban were asked to participate in the study. Face –to- face in-depth interviews were conducted with 8 consenting refugee women.

In this paper we are able to document a number of challenges faced by refugee women in Durban, KwaZulu-Natal regarding medical xenophobia and how these impact negatively on their reproductive health.

### Problem statement

The number of refugee women in South Africa continues to grow as the number of refugees and foreigners entering South Africa is increasing. ^[7]^ Among them are women from the Great Lakes region, whom due to the war situation in their countries have been exposed to health risks from their home, along the way to their destination, and carried those risks into the host country. Upon their arrival in South Africa, the refugee women do not undergo any screening, yet many of them are survivors of rape and many other acts of sexual violence. There is paucity of literature on the experiences of refugee women regarding access to reproductive health services in South Africa. This lack of documentation is a threat, not only to the refugee community, but also to the host, because after integration in the host community, there are inter-marriages and cohabitation between local people and refugees. Furthermore, the latter are continuously reproducing despite the socio-economic life conditions that they are living under.

### Aim of the study

The purpose of this observational study was to describe the voices of women refugees and their challenges regarding reproductive health services in public health institutions in Durban, KwaZulu Natal

## Research methodology

### Study design

This was an exploratory qualitative study using interpretivist paradigm. Thematic content analysis guided this qualitative study. Participants were requested to participate in the study if they were from one of three Great Lakes countries, that is the DRC, Burundi and Rwanda and living in Durban as refugees. Women between ages 18 and 49, and having sought reproductive health services at one of the public health facilities situated in the city of Durban were asked to participate.

### Setting

The study was conducted in eThekwini district, Durban KwaZulu-Natal. Interviews were conducted in two churches that host most refugees from the Great Lakes countries. Interviews were held in the private space within the churches at times suitable for participants.

### Sampling

Purposive sampling was used to recruit participants in this study as it allowed the researcher to select participants who were able to provide rich information about the phenomenon that was being studied. ^[8]^ Refugee women who originated from the Great Lakes region (Rwanda, Burundi and DRC), who live in Durban, KwaZulu-Natal and have sought reproductive health services in any public healthcare facilities in Durban were approached to participate in the study.

### Data collection

Data was collected in September 2017 using a semi-structured interview schedule. In-depth individual interviews were done with 8 participants until data saturation was reached. Before each interview started the researcher went through the informed consent and information leaflet with participants after which they signed and then proceeded to interviews. All interviews were audio-recorded and the researcher took notes during interviews. The interviews took place in three selected church boardrooms in Durban KwaZulu-Natal at suitable times for each participant. Each interview took 30 to 45 minutes.

### Data analysis

Data analysis was done after each interview to allow the researcher to monitor data saturation. The in-depth interviews were transcribed verbatim. The researcher started by transcribing the recorded data at the same time paying attention to the notes taken during each interview with the participants, to ensure no information is left behind. Then the researcher read and re-read the transcribed data, highlighting, identifying and organising the main categories with similar ideas, which she later grouped into themes and sub-themes to help her outline the analysis before the actual analysis took place. The researcher then proceeded by summarising and interpreting the findings and themes.

### Trustworthiness

To establish the trustworthiness of the qualitative data, the principles discussed by Lincoln and Guba ^[9]^ were used. To ensure credibility of the study, notes were taken during face-to-face interviews and transcribed after each interview. The interviews were audio-recorded, then transcribed verbatim to make sure that all participants’ responses are captured appropriately. Following the transcription of the recorded interviews, the transcripts were returned to the study participants who confirmed the accuracy of information they provided during data collection. The interviews were conducted in a quiet environment to avoid sounds that might affect participants’ voice recording.

### Ethical considerations

Permission to conduct the study was obtained from the Humanities and Social Sciences Research Ethics Committee of KwaZulu-Natal, reference number: HSS/0224/016M. Permission was obtained from the 2 churches where participants were recruited. Participants were given an information letter on the study, and they willingly signed an informed consent. Confidentiality and anonymity of the data were ensured by the researcher.

## Results

It is evident from Table 1 that all study participants had a refugee status in South Africa Two major themes emerged from the interviews and included negative experiences or challenges and the positive experiences. The themes and sub-themes are displayed under table 2.

**Table 1:** Demographic characteristics of participants

| **Pseudonym** | **Age** | **Level of education** | **Country of origin** | **No. of children** | **Marital status** | **Type of document** | **Occupation** |
| --- | --- | --- | --- | --- | --- | --- | --- |
| Participant 1 | 43 | Diploma | Rwanda | 3 | Married | Refugee status | Unemployed |
| Participant 2 | 48 | Diploma in teaching | Rwanda | 4 | Married | Refugee status | Social worker |
| Participant 3 | 25 | Matric | Burundi | 2 | Single | Refugee status | Unemployed |
| Participant 4 | 40 | 2 years secondary school | Burundi | 2 | Married | Refugee status | Small business |
| Participant 5 | 36 | Diploma | Rwanda | 3 | Married | Refugee status | School librarian assistant |
| Participant 6 | 26 | Diploma | DRC | 1 | Married | Refugee status | Housewife |
| Participant 7 | 24 | Matric | Rwanda | None | Single | Refugee status | Student |
| Participant 8 | 47 | Diploma | DRC | 3 | Married | Refugee status | Housewife |

**Table 2:** Themes and sub-themes

| Themes | Sub-themes |
| --- | --- |
| **1. Negative experiences/challenges** | 1.1. Medical xenophobia<br>1.2. Language barrier<br>1.3. Discrimination<br>1.4. Unprofessionalism<br>1.4.1. Lack of health education and customer care training<br>1.4.2 Failure to obtain consent and lack of confidentiality<br>1.4.3. Ill-treatment<br>1.5. Financial challenges<br>1.6. Internalised fear<br>1.7. Shortage of health personnel and overcrowding of public hospitals<br>1.8. Religious and cultural hegemony |
| **2. Positive experiences** | 2.1. Positive treatment and care<br>2.2. Social support |

### Theme 1: Medical xenophobia and discrimination

Six out of eight participants experienced xenophobia during their first encounter with health services

> She stood up from her chair shouting and pointing fingers at me saying “you foreigners you are using our hospitals but you do not pay tax and you like to have babies”. I didn’t know what to say so I walked out of the room and she followed me saying that I do not want to listen when she is telling me what to do. I was afraid so I left that clinic and never went back” [said in an angry tone]. **Participant 4**

Language was seen as a barrier between participants and clinic staff and believed by participants to be another factor contributing to medical xenophobia

> “When you just start saying hello, they [nurses]can’t even pay attention to you because you are speaking English but if you are speaking in IsiZulu they will talk to you nicely”. Participant 6

The local people use critical stereotypes against African foreigners such as labelling them as “makwerekwere”. Refugee women viewed that as being discriminated in front of all other patients seeking reproductive health services

> “The day I went to give birth it was the worst day of my life, there was no one to take care of me. I was calling the nurse telling them that I feel like pushing and I was feeling the baby’s head coming out but no one came” …. “You kwerekwere you always like to scream just keep quiet you are still far….” **Participant 5**

Almost all the participants encountered some form of discrimination at the public hospital due to being foreigners in the country. One participant described one such experience:

> “The first time when I went to the hospital, I took my child for immunisation, the nurses looked at us like we are not people like them. They started swearing at us saying we come here to get babies, that we don’t pay anything and we use their tax money, then they continued saying we must stop having babies.” **Participant 1**

Participants felt that discrimination is so widespread that it gives rise to a sense of not belonging and a feeling that being a foreigner means that one is not a human being.

> “You can’t be treated the same as the locals if you want to access these services”. **Participant 3**
>
> “The nurses don’t treat us like human being; the challenge is to go to the hospital when you are a foreigner because the treatment we get is very bad. To me being a refugee is a big problem when it comes to accessing health services.” **Participant 6**

### Theme 2: Language barrier

Language barrier is the inability to effectively communicate with others, which may lead to medical errors by impending patient- provider communication. ^[3]^ Six (6) out of eight (8) participants faced the challenge of the language barrier in public hospitals because they could not speak the local isiZulu language and this had a negative effect on the nature of services they received at these hospitals.

> “It was quite difficult, I couldn’t communicate. I used to take someone along with me to help me translate. It was still difficult because the translator was also a refugee and her English was not good but she was better than me.” **Participant 3**
>
> so, the big challenge was that I couldn’t speak isiZulu, others were shouting in their own language”. **Participant 5**

There is a strong relationship between language and power, with those exercising power in society using it as a tool to oppress the powerless. In this instance, nurses hold power as they are in a position of authority, and they abuse their power through language to oppress powerless refugee women. There is a stereotype amongst black South Africans, in this case IsiZulu speaking people that every black person is an isiZulu speaker, and this gives rise to cultural imperialism in which a dominant group wants to impose its norms onto another group of people, in this instance foreigner

> “Why are you not talking Zulu why you don’t talk my language and you look like me? We’re all Africans but you can’t talk my language”. **Participant 8**
>
> “We were so many in the clinic and the nurse spoke to me in Zulu; when I tried to tell her that I did not understand asking her to explain in English and she shouted saying, “You people go back to your country we are tired of you… she continued, saying that she has to talk in Zulu because English is not their language.” **Participant 6**

### Theme 3: Lack of professionalism

All the participants encountered unprofessional behaviour from the nurses in the public hospitals and clinics. This was seen at various levels, and various sub-themes emerged from it which were

- Lack of health education and customer care training,
- Failure to obtain consent and lack of confidentiality,
- Ill-treatment.:

#### 3.1: Lack of health education and customer care training

All the participants complained about the lack of health education. When participants visited a family planning clinic to request a family planning method, they expected to be educated on the methods to make informed choices

> “So, they [nurses] want to force us to take injections or tablets without even explaining to us which one is better.” **Participant 1**
>
> I went having a bit of knowledge about it but the first thing the lady [a nurse] wanted to do to me was to inject me and I quickly rejected that [bewildered look]. I wanted tablets and the nurse was not happy then she just gave me the tablets without explaining anything so I had to rely on other people outside. **“Participant 2**

#### 3.2: Failure to obtain informed consent

Participants were required to partake in procedures that they did not wholly agree on, which was a gross violation of the rights of patients and lack of professionalism by the healthcare providers concerned. Patricia described how she was tricked into signing a consent form even though she did not know the reason why she was signing it in the first place:

> “The c/section was not a good experience in government hospital, it was something forced onto me and I was tricked to signing consent but they did not explain that I was going to have an operation.” **Participant 3**
>
> “They force you yeah … they force you to take injection.” **Participant 8**

#### 3.3: Ill -treatment

All the participants mentioned that they were ill- treated at some point in public hos*pitals*,

> “The nurses they look at us like we are not people like them they start swearing at us. They are so rude.” **Participant 1**
>
> “…the majority of nurses are very bad”. **Participant 6**

### Theme 4: Internalised fear

Participants reported fear emanating from the fact that they were foreigners in South Africa and the negative treatment they continually received in public hospitals and in other public spheres

> “So, they use their language because they think you don’t understand what they are saying but when you read their body language you can figure out what they are talking about then it makes you scared to go there. Even when we tell our friends they are also scared to go the hospital when they are pregnant, they like to go when they are about six or seven months.” **Participant 1**

## Discussion

The aim of this paper was to describe the voices of women refugees and their challenges regarding reproductive health services in public health institutions in Durban, KwaZulu Natal. Refugees face discrimination and medical xenophobia when seeking healthcare services at public health facilities in South Africa. ^[2, 3]^ In this study, the findings revealed that all participants experienced medical xenophobia and discrimination in public healthcare facilities while seeking reproductive health services. Participants reported facing negative attitudes from healthcare professionals, mostly nurses. Participants also expressed feelings of being neglected and others denied treatment by nurses simply because they were foreigners.

Refugee women who participated in the study faced challenges of language barriers and bad attitudes from the healthcare workers in public hospitals and clinics when seeking reproductive health services. Communication between health professionals and the participants was a challenge due to language differences. Nurses insisted on speaking isiZulu to the foreigners who could hardly understand a word of it. The study findings also revealed that, while seeking the reproductive health services in the public sectors, women who were able to speak isiZulu were assisted professionally, as opposed to women refugees and other foreign nationals, who could not explain themselves in isiZulu.

Women reported that they have no interpreters at public health facilities and that they struggle to communicate with the healthcare professionals. These findings were supported by Apalata, Kibiribiri ^[10]^, in their study which revealed that refugee women face many negative experiences with reproductive health services, not only because they cannot communicate but because there are no interpreters or facilities to convey their messages. Consequently, refugee women end up either returning home untreated or having to bring their children or friends who are also not fluent in the local languages to assist with interpretation. However, literature shows that when interpretation is done by an untrained person, there are risks to transmit a wrong message resulting in wrong diagnosis and treatment which later affect the client negatively. ^[11]^

The number of refugees and foreigners entering South Africa is increasing and South Africans will have to be educated concerning refugees and their wellbeing. ^[7]^ Lack of health education and lack of good customer-care training for healthcare workers were seen as being unprofessional. Lack of understanding of the refugees“ situation and their rights to healthcare services and limited or non-existent good customer-care behavior affect refugees“ healthcare access and usage. The findings revealed that nurses in the public sectors lack training in how to treat or deal with vulnerable people, in this case women refugees in the host country. The focus should be on improving the management and care of refugees.^[12]^ Refugees reported negative experiences which they associated with negligence by healthcare professionals, ignorance, lack of education about refugees“ background and very limited customer-care training.

The study highlighted challenges regarding healthcare provider’s failure to get consent from clients before doing medical procedures and giving medication. Some refugee women reported being deceived and forced into having medical procedures, such as caesarean section and HIV testing, without their consents. Other participants reported being forced to have an injection for family planning against their will. Refugee women also felt that confidentiality of their information was compromised. Apalata, Kibiribiri ^[10]^ also states challenges of lack of consent and confidentiality, and refugee women being forced to deliver by caesarean without their consent

Feelings of rejection and ill-treatment by refugees when attending public healthcare institutions has resulted in the majority no longer interested in attending public hospitals. ^[10]^ In this study, it was noted that refugee women were ill-treated by healthcare providers in the public sector. All the participants reported that they have been ill-treated at some point in public hospitals and clinics, especially by nurses. Some participants reported that they would never have any more babies in South Africa due to the ill-treatment they received while attending antenatal clinics, during pregnancy and after delivery. While the above are the results of ill-treatment of the refugees at public health facilities ^[13,14]^ it had created an internalized fear in the refugee community. Many of them as reported in this study, end up visiting private doctors even though they do not have money to afford their services, but they still feel confident and trust the private sector than public

## Limitations

The study focused on one sub-district and this limits generalisability of findings. However, they align with previously published studies. Non-church goers were not included in the study and that limited the views of refugee women on the subject of inquiry.

## Recommendations

This study can be replicated in other parts of Southern Africa region where refugees are hosted to document women experiences with sexual and reproductive health services. There is also a need to conduct research that looks at alternative ways used by refugee women when they fail to access reproductive health care services at public health facilities. Further research to explore the risk factors associated with refugee women lack of reproductive health services in this era of HIV is needed. The refugee community churches and other non-religious institutions should partner with government and non-governmental organisations active in the area of refugees to decrease the challenges faced by refugee women

## Conclusion

In conclusion, this study described the voices of women refugees and their challenges regarding reproductive health services in public health institutions in Durban, KwaZulu Natal. Refugee women continue to face challenges when seeking reproductive health services in public health facilities. They remain vulnerable when they are denied access to reproductive health services and they face numerous challenges including contracting HIV and other sexually transmitted infections. They also face complications during pregnancy and post-delivery and this poses a threat to public health.

## Acknowledgements

The authors acknowledge the University of KwaZulu-Natal, the School of Nursing and Public Health for providing resources to complete the study.

## Competing interests

The authors declare that they have no competing interests

## Authors’ contributions

Y.M conducted the study for a masters degree and prepared the manuscript. EMM supervised the study, provided guidance in preparation and completion of the manuscript

